# Structure of KCNH2 cyclic nucleotide-binding homology domain reveals a functionally vital salt-bridge

**DOI:** 10.1101/790154

**Authors:** Ariel Ben-Bassat, Moshe Giladi, Yoni Haitin

## Abstract

Human KCNH2 (hKCNH2, Ether-à-go-go (EAG)-Related Gene, hERG) are best known for their role in cardiac action potentials repolarization and have key roles in various pathologies. As other KCNH family members, hKCNH2 contains a unique intracellular complex crucial for channel function, consisting of an N-terminal eag domain and a C-terminal cyclic nucleotide-binding homology domain (CNBHD). Previous studies demonstrated that the CNBHD is occupied by an intrinsic ligand motif (ILM), in a self-liganded conformation, providing a structural mechanism for the lack of KCNH channels regulation by cyclic nucleotides. While significant advancements in structural and functional characterizations of the CNBHD of KCNH channels have been made, a high-resolution structure of the hKCNH2 intracellular complex was missing. Here, we report the 1.5 Å resolution structure of the hKCNH2 channel CNBHD. The structure reveals the canonical fold shared by other KCNH family members, where the spatial organization of the ILM is preserved within the β-roll region. Moreover, measurements of small-angle X-ray scattering profile in solution, as well as comparison with a recent nuclear magnetic resonance (NMR) analysis of hKCNH2, revealed high agreement with the structure, indicating an overall low flexibility in solution. Importantly, we identified a novel salt-bridge (E807-R863), which was not previously resolved in the NMR and cryogenic electron microscopy (cryo-EM) structures. Strikingly, electrophysiological analysis of charge reversal mutations revealed its crucial role for hKCNH2 function. Moreover, comparison with other KCNH members revealed the structural conservation of this salt-bridge, consistent with its functional significance. Together with the available structure of the mouse KCNH1 intracellular complex, and previous electrophysiological and spectroscopic studies of KCNH family members, we propose that this salt-bridge serves as a strategically positioned linchpin to support both the spatial organization of the ILM and the maintenance of the intracellular complex interface.

**Summary:** Human KCNH2 are key channels governing cardiac repolarization. Here, a 1.5 Å resolution structure of their cyclic nucleotide-binding homology domain is presented. Structural analysis and electrophysiological validation reveal a novel salt-bridge, playing an important role in hKCNH2 functional regulation.

## Introduction

hKCNH2 (the human Ether-à-go-go (EAG)-Related Gene), also known as hERG1, is a member of the KCNH voltage-dependent delayed rectifier potassium channel family. Best known for their critical role in cardiac action potentials repolarization (Sanguinetti & Jurkiewicz, 1990; Sanguinetti *et al.*, 1995; Curran *et al.*, 1995; Trudeau *et al.*, 1995), these channels are important regulators of cellular excitability (Warmke *et al.*, 1991; Ganetzky *et al.*, 1999). Moreover, hKCNH2 have key roles in cardiac pathologies (Sanguinetti & Tristani-Firouzi, 2006) as well as neuronal and mental diseases such as epilepsy (Zhang *et al.*, 2010) and schizophrenia (Huffaker *et al.*, 2009). Importantly, these channels are notoriously sensitive to adverse drugs interactions, resulting in drug-induced Long QT syndrome (LQTS) (Sanguinetti & Tristani-Firouzi, 2006), a potentially lethal cardiac disorder, and are known to harbor numerous LQTS-related mutations (Curran *et al.*, 1995). Finally, similar to other KCNH family members, hKCNH2 expression was demonstrated in various tumors (Bianchi *et al.*, 1998), possibly indicating additional non-canonical pathophysiological roles.

Comprising of four identical subunits surrounding a centrally located ion conductive pore, each hKCNH2 subunit demonstrates the characteristic voltage-dependent potassium channel topology, consisting of six membrane-spanning domains, with cytoplasmic N- and C-termini (Warmke *et al.*, 1991). However, unique to the KCNH channels family, the N-terminal region contains an eag domain and the C-terminal region contains a cyclic nucleotide-binding homology domain (CNBHD), with both domains forming an intracellular complex crucial for channel function (Morais Cabral *et al.*, 1998; Gustina & Trudeau, 2009; Haitin *et al.*, 2013). Importantly, it has recently become clear that many of the specialized gating properties of KCNH channels arise from their structurally distinct intracellular domains (Morais-Cabral & Robertson, 2015). Indeed, many LQTS-related mutations are scattered throughout the intracellular complex, suggesting it plays critical roles during protein biogenesis, trafficking and function (Chen *et al.*, 1999; Anderson *et al.*, 2006; Harley *et al.*, 2012).

While the CNBHD of KCNH channels shows a high level of structural conservation with the cyclic nucleotide binding domain (CNBD) of hyperpolarization-activated cyclic nucleotide–gated (HCN) channels, it does not bind cyclic nucleotides (Robertson *et al.*, 1996; Brelidze *et al.*, 2009). Instead, recent molecular structures, of multiple KCNH homologues, revealed that the cyclic nucleotide binding pocket within the CNBHD is occupied by a short β-strand fold, termed the ‘intrinsic ligand’ motif (ILM), perfectly mimicking the occupancy of cyclic nucleotides within the CNBD of HCN channels (Brelidze *et al.*, 2012; Marques-Carvalho *et al.*, 2012; Haitin *et al.*, 2013). Critically, the discovery that the CNBHD of KCNH channels is in a self-liganded conformation was not predicted by mere sequence analyses and resulted directly from structural studies of this domain. Moreover, this short three residue stretch was recently shown to be involved in shaping the gating properties of KCNH1 channels (Zhao *et al.*, 2017), suggesting a mechanistic role for the otherwise functionally orphan CNBHD.

The recent ‘resolution revolution’ of single particle cryogenic electron microscopy (cryo-EM) did not skip the KCNH channels family (Whicher *et al.*, 2016; Wang & MacKinnon, 2017). However, while providing groundbreaking insights into the architecture of the intact channel, the resolution ranges between ~4.5 – 5.5 Å at the cytosolic eag domain and CNBHD. Thus, the derivation of atomic resolution details regarding the organization of the ILM is limited. Here, we used size-exclusion chromatography combined with multi-angle light scattering (SEC-MALS), X-ray crystallography and small angle X-ray scattering (SAXS) to characterize the structural properties of the CNBHD of hKCNH2. Our results provide the first high-resolution crystal structure of hKCNH2, resolved at 1.5 Å resolution. In agreement with the available structural information, the structure reveals that the CNBHD of hKCNH2 shares a similar fold with other members of the family. Accordingly, the spatial organization of the ILM is preserved within the canonical β-roll region (Brelidze *et al.*, 2012, 2013; Marques-Carvalho *et al.*, 2012). Intriguingly, comparison of our X-ray structure with a previously determined NMR ensemble (Li *et al.*, 2016) suggests the presence of highly labile regions within this domain. However, followed by ensemble optimization method (EOM) analysis, SAXS data shows good agreement between the solution properties and the crystal structure, indicating the CNBHD of hKCNH2 is overall highly rigid. Finally, the crystal structure revealed the presence of a previously overlooked, strategically located, salt-bridge stabilizing the ILM. Importantly, electrophysiological analyses of charge reversal mutants revealed its crucial role for normal channel function.

## Materials and methods

### Protein expression and purification

The CNBHD (residues 734–864) of human KCNH2 (hKCNH2; Genebank accession code NM_000238.3) was sub-cloned using 5′ NcoI and 3′ HindIII sites into a pETM11 vector containing an N-terminal hexa-histidine affinity tag followed by a tobacco etch virus protease (TEV) cleavage site and a GAMG cloning sequence. The construct was verified by sequencing. Proteins were overexpressed and purified as previously described (Haitin *et al.*, 2013), using immobilized metal affinity chromatography and size-exclusion chromatography. Briefly, *E. coli* T7 express competent cells (New England Biolabs, Ipswich, MA, USA), transformed with the CNBHD construct, were grown in Terrific Broth medium at 37°C until reaching OD_600nm_ = 0.6 and induced at 16°C by adding 0.25 mM isopropyl β- D-1-thiogalactopyranoside (IPTG; Formedium, Norfolk, UK). Proteins were expressed at 16°C for 16-20 h and harvested by centrifugation (10,000 x g for 10 min). Cells were re-suspended in buffer A, containing (in mM) 150 NaCl, 50 Bis-Tris, pH 6.5, and 1 tris(2-carboxyethyl)phosphine (TCEP), supplemented with 15 mM imidazole, 2.5 μg ml^−1^ DNaseI, 5 mM MgCl_2_, 10 mg lysozyme, 1 mM phenylmethane sulfonyl fluoride (PMSF) and Protease Inhibitor Cocktail Set III (Calbiochem, San Diego, CA, USA). Cells were lysed with an EmulsiFlex C-5 homogenizer (Avestin, Ottawa, ON, Canada) and the lysate was cleared by centrifugation at 32,000 x g for 45 minutes at 4 °C. The supernatant was then loaded onto a Ni^2+^ affinity resin column (HisTrap HP, GE Healthcare, Chicago, IL, USA) followed by washing with buffer A containing 27 mM imidazole and eluted using buffer A supplemented with 300 mM imidazole. The hexa-histidine tag was removed by TEV cleavage overnight at 4 °C. Imidazole was removed using a HiPrep 26/10 desalting column (GE Healthcare, Chicago, IL, USA), equilibrated with buffer A containing 15 mM imidazole. The cleaved protein was loaded onto a second Ni^2+^ column with 15 mM imidazole to remove the cleaved hexa-histidine tag and TEV protease. The flow-through was collected, concentrated to 4–5 mL, and loaded onto a HiLoad 16/60 Superdex 75 column (GE Healthcare, Chicago, IL, USA), equilibrated with gel filtration buffer, containing (in mM) 150 NaCl, 5 dithiothreitol (DTT) and 20 Bis-Tris, pH 6.5. Finally, pooled fractions were concentrated to ~220 μM (3.3 mg/ml) using a 3 kD molecular weight cut-off concentrator (EMD-Millipore, Burlington, MA, USA), flash-frozen in liquid N_2_ and stored at −80 °C until use.

### Size-exclusion chromatography multi-angle light scattering (SEC-MALS) analysis

Experiments were performed on an HPLC system (Shimadu, Kyoto, Japan), equipped with autosampler and UV detector, using a preequilibrated analytical SEC column (Superdex 200 10/300 GL; GE Healthcare, Chicago, IL, USA) with gel filtration buffer. A Sample of the purified CNBHD (2 mg/ml, 50 μl) was loaded onto the column and analyzed using an inline 8-angle light-scattering detector, followed by a differential refractive-index detector (Wyatt Technology, Santa Barbara, CA, USA). Refractive index and MALS readings were analyzed using the Astra software package (Wyatt Technology) to determine the molecular mass.

### Protein crystallization and data collection

Crystals of the CNBHD were grown at 19 °C using sitting-drop vapor diffusion by mixing a 1:1 ratio (v/v; Mosquito, TTP Labtech, Hertfordshire, UK) of protein solution at 200 μM and a reservoir solution, containing 20% (w/v) PEG 3350, 2% (v/v) Tacsimate, 0.1M Bis-Tris, pH 6.8. This condition produced crystals within two days, which grew to a maximum size of about 200 × 100 × 100 μm after 7 days. For diffraction data collection, crystals were immersed in liquid N_2_ after cryoprotection with 20% glycerol. Data were collected at 100 °K on beamline ID23-1 of the European Synchrotron Radiation Facility (ESRF; Grenoble, France), using a wavelength of 0.972 Å. Integration, scaling and merging of the diffraction data were performed using the XDS program (Collaborative Computational Project, 1994). The crystals belonged to space group C222_1_ (a = 57.49 Å, b = 90.65 Å and c = 61.11 Å with α = β = γ = 90°).

### Structure determination

The structure was solved by molecular replacement using the programs PHASER (McCoy, 2006) and PHENIX (Adams *et al.*, 2010) using a modified chain A of the KCNH1 intracellular complex structure (Protein Data Bank code 4LLO) as the search model. Data collection and refinement statistics are summarized in Table 1. Each asymmetric unit contained a single protein chain. Iterative model building and refinement were carried out in PHENIX with manual adjustments using COOT (Emsley & Cowtan, 2004). All structural illustrations were prepared with UCSF Chimera (https://www.cgl.ucsf.edu/chimera).

**Table 1.**
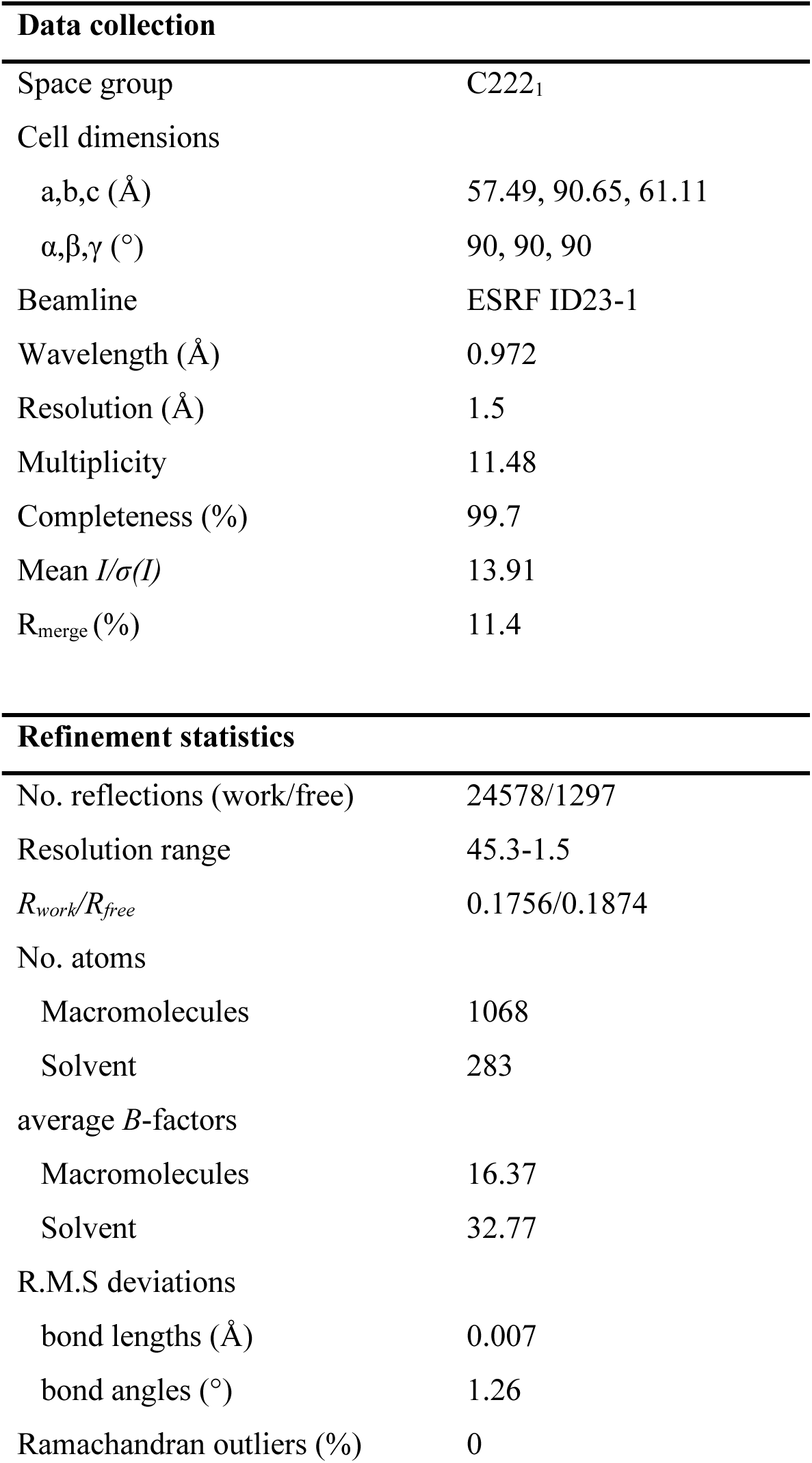
Crystallographic statistics

| **Data collection** |  |
| --- | --- |
| Space group | C222<sub>1</sub> |
| Cell dimensions |  |
| a,b,c (Å) | 57.49, 90.65, 61.11 |
| $\alpha,\beta,\gamma$ (°) | 90, 90, 90 |
| Beamline | ESRF ID23-1 |
| Wavelength (Å) | 0.972 |
| Resolution (Å) | 1.5 |
| Multiplicity | 11.48 |
| Completeness (%) | 99.7 |
| Mean $I/\sigma(I)$ | 13.91 |
| R<sub>merge</sub> (%) | 11.4 |
| **Refinement statistics** |  |
| No. reflections (work/free) | 24578/1297 |
| Resolution range | 45.3-1.5 |
| $R_{work}/R_{free}$ | 0.1756/0.1874 |
| No. atoms |  |
| Macromolecules | 1068 |
| Solvent | 283 |
| average $B$-factors | |
| Macromolecules | 16.37 |
| Solvent | 32.77 |
| R.M.S deviations |  |
| bond lengths (Å) | 0.007 |
| bond angles (°) | 1.26 |
| Ramachandran outliers (%) | 0 |

### SAXS Data Collection and Analysis

SAXS data were measured at beamline BM29 of the ESRF (Grenoble, France). Data were collected at 5 °C with X-ray beam at wavelength λ = 1.0 Å, and the distance from the sample to detector (PILATUS 1 M, Dectris Ltd) was 2.867 meters, covering a scattering vector range (*q* = 4 πsinθ/λ) from 0.0025 to 0.5 Å^−1^. 10 frames of two-dimensional images were recorded for each buffer or sample, with an exposure time of 1 s per frame. The 2D images were reduced to one-dimensional scattering profiles and the scattering of the buffer was subtracted from the sample profile using the software on site. Samples were analyzed in gel filtration buffer. To account for possible inter-particle effects, each sample was measured at three concentrations (Table 2). The lowest concentration curve was merged with a higher concentration curve at *q* ~ 0.2 Å^−1^ to prevent distortion of the low-angle data while preserving high signal-to-noise ratio at the higher angles, which are far less sensitive to interparticle effects (Koch *et al.*, 2003). The experimental radius of gyration (*R*_g_) was calculated from data at low *q* values in the range of qRg < 1.3, according to the Guinier approximation: ln*I*(*q*) ≈ ln(*I*_0_) − *R*_g_2*q*2/3 using PRIMUS (Konarev *et al.*, 2003). The Porod volume was derived from the paired-distance distribution function (PDDF or *P*(r)) calculated using GNOM (Petoukhov *et al.*, 2012). The solution scattering of the crystal structure was calculated using CRYSOL (Svergun *et al.*, 1995).

**Table 2.** Data collection and scattering-derived parameters.

| **Data collection parameters** |  |
| --- | --- |
| Sample | hKCNH2 734-864 |
| Beamline | ESRF BM29 |
| Beam geometry (mm<sup>2</sup>) | 0.7 x 0.7 |
| Wavelength (Å) | 1.0 |
| Q range (Å<sup>-1</sup>) | 0.0025 – 0.5 |
| No. of frames | 10 |
| Exposure per frame (seconds) | 0.5 |
| Concentration (mg/ml) | 3.0, 1.5, 0.7 |
| Temperature (°C) | 5 |
| Structural parameters |  |
| $I_0$ (from Guinier) | $11.3 \pm 0.1$ |
| $R_g$ (Å) (from Guinier) | $15.9 \pm 0.2$ |
| Porod volume (10<sup>3</sup> Å<sup>3</sup>) | 27.23 |
| $\chi^2$ CRY SOL | 1.4 |
| $\chi^2$ EOM | 0.8 |
| Software employed |  |
| Primary data reduction | AUTOMAR |
| Data processing | PRIMUS |
| Modelling | CRY SOL, EOM |

<sup>1</sup> ± S.E. <sup>2</sup> ± 10% (estimated range).

### Ensemble optimization method (EOM) analysis

The software RANCH (Bernadó *et al.*, 2007) was used to generate a pool of 10,000 stereochemically feasible structures with a random linker between the α’F and αA. This pool was used as input for GAJOE (Bernadó *et al.*, 2007) that selects an ensemble with the best fit to the experimental data using a genetic algorithm: 50 ensembles of 20 orientations each were “crossed” and “mutated” for 1,000 generations and the process was repeated 50 times.

### Two electrode voltage clamp

Oocytes were obtained by surgically removal of ovary pieces from *Xenopus laevis* frogs (Xenopus 1, MI, USA), anesthetized with 0.15% tricaine (Sigma-Aldrich). The procedures for surgery and maintenance of frogs were approved by the animal research ethics committee of Tel Aviv University and in accordance with the Guide for the Care and Use of Laboratory Animals (National Academy of Sciences, Washington DC). Oocytes were de-folliculated using 1 mg/ml collagenase (type IA, Sigma-Aldrich) in Ca^2+^-free ND96 solution, containing (in mM): 96 NaCl, 2 KCl, 1 MgCl_2_, and 5 HEPES, pH = 7.5 (ND96), for about 1 h. Stage V and VI oocytes were selected for DNA injection and maintained at 18°C in ND96, supplemented with 1 mM pyruvate and 50 μg/ml gentamycin. For channel expression, intranuclear injections of 100 ng/μl hKCNH2 cDNA (in pcDNA3 vector; 10 nl per oocyte) were performed using a Nanoject injector (Drummond, PA, USA). Standard two-electrode voltage-clamp measurements were performed at room temperature (22–24°C) 2-5 d after DNA microinjections. Oocytes were placed into a 100-μl recording chamber and perfused with a modified ND96 solution (containing 0.1 mM CaCl_2_). Whole-cell currents were recorded using a GeneClamp 500 amplifier through the pCLAMP 9.2.1.9 software (Axon Instruments, Inc.). Glass microelectrodes (A-M systems, Inc.) were filled with 3M KCl and had tip resistances of 0.2-1 MΩ. Current signals were digitized at 1 kHz and low pass filtered at 0.2 kHz. The holding potential was −80 mV. Leak subtraction was performed off-line, using steps from −120 to −90 mV, assuming that the channels are closed at −80 mV and below. Errors introduced by the series resistance of the oocytes were not corrected and were minimized by keeping expression of the currents below 10 μA. The steady-state voltage dependence of activation (G-V) was measured by plotting the tail current amplitude versus the previous test pulse voltage and fit with a Boltzmann function:

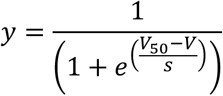

where V_50_ is the half-maximal activation potential and s is the slope factor. All data are presented as mean ± SEM, and n represents the number of cells. Current deactivation was fit with a double-exponential function:

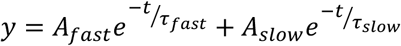

where *t* is time and *τ* is the time constant of deactivation.

Statistical analyses were performed using a student’s t-test. P < 0.05 was considered statistically significant.

## Results

### Structural characterization of the CNBHD of hKCNH2

In order to elucidate the molecular organization of the CNBHD of hKCNH2, we first designed a soluble construct, amiable for downstream purification and crystallization. Specifically, similar to the strategy previously used for structural studies of the CNBHD of mouse KCNH1 (Marques-Carvalho *et al.*, 2012; Haitin *et al.*, 2013), we truncated most of the C-linker, connecting the S6 and the CNBHD (Marques-Carvalho *et al.*, 2012), and terminated the construct just downstream of the ILM (Brelidze *et al.*, 2012). This construct, termed CNBHD hereafter, spans residues 734-864 and consists of a remaining C-linker segment (734-747) and the full-length CNBHD (748-864) (Fig. 1A).

**Figure 1.**
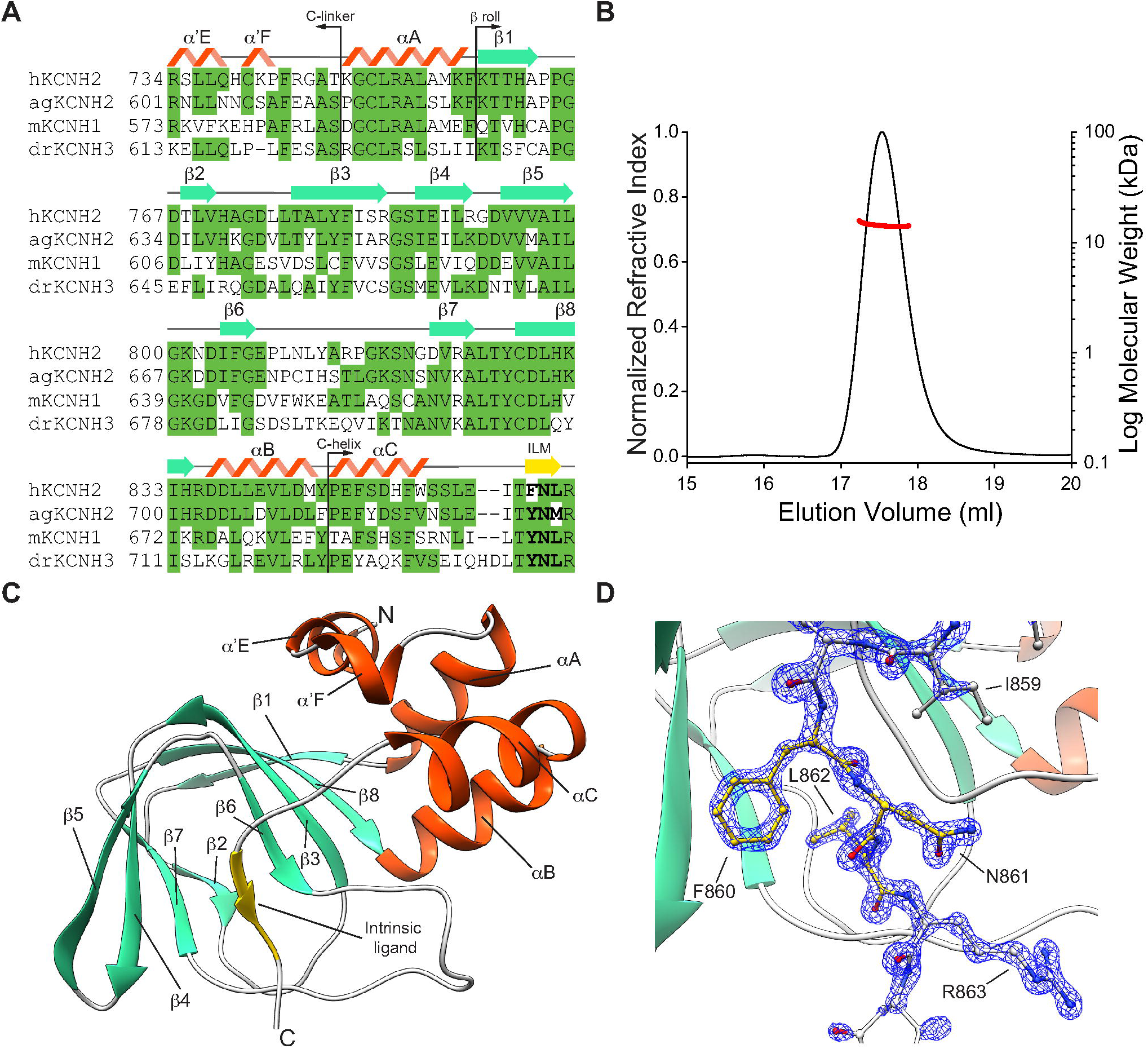
Structure of hKCNH2 CNBHD. **(A)** Sequence alignment of the CNBHD from various KCNH family members whose structures are available. Secondary structure of hKCNH2 is indicated above the sequence. Loops and non-helical secondary structure are marked as solid gray lines. Residues conserved in KCNH channels are shown in green. The intrinsic ligand motif (ILM) is bolded. **(B)** SEC-MALS analysis of the purified hKCNH2 CNBHD. The black curve represents the normalized refractive index and the red curve indicates the calculated molecular weight in kDa. **(C)** Structure of hKCNH2 CNBHD in cartoon representation, colored according to its secondary structure. (D) A detailed illustration of the ILM (yellow) and flanking residues (grey) shown as sticks and colored by element, overlaid with 2Fo-Fc map contoured at 3σ (blue mesh).

Next, using SEC-MALS, we determined that the purified CNBHD is monodispersed with a molecular mass of 14.3 ± 0.6 kDa (Fig. 1B), close to the calculated mass of the protein (14,929.22 Da). Thus, in the absence of a C-linker, the CNBHD of hKCNH2 is monomeric in solution, similar to the observed stoichiometry of purified CNBHDs from other members of the KCNH family (Brelidze *et al.*, 2012, 2013; Marques-Carvalho *et al.*, 2012).

Following purification and solution characterization, we proceeded with CNBHD crystallization. The CNBHD of hKCNH2 readily crystallized in the space group C222_1_ with a single molecule in the asymmetric unit and diffracted X-rays to 1.5 Å resolution (Table 1). The structure was solved by molecular replacement, using the CNBHD of mouse KCNH1 intracellular complex (Haitin *et al.*, 2013) as a search model. The structure of the CNBHD of hKCNH2 (Fig. 1C) can be subdivided into four distinct segments: (i) The remnant C-linker, composed of helices α’E and α’F, organized in a helix-turn-helix configuration; (ii) The β-roll, featuring the canonical eight β strands fold (β1 – β8), mediating ligand binding in HCN and cyclic nucleotide-gated ion channels (Craven & Zagotta, 2006); (*iii*) The C-helix (helix αC), which, together with αB, flanks the β-roll and was shown to underlie ligand-induced gating enhancement of cyclic nucleotide-gated channels (Puljung & Zagotta, 2013); And, (*iv*) the KCNH family unique ILM, consisting of a short β strand (β9) (Fig. 1D), which is involved in the formation of the interaction interface with the N-terminal eag domain (Haitin *et al.*, 2013; Wang & MacKinnon, 2017), and differentially affects KCNH channels activation (Zhao *et al.*, 2017) and deactivation (Codding & Trudeau, 2019). Together, the overall fold of the CNBHD of hKCNH2 is reminiscent to that observed in other CNBD-containing proteins (Rehmann *et al.*, 2007).

### The hKCNH2 CNBHD X-ray and solution NMR structures differ

Recently, the solution structure of the CNBHD of hKCNH2 was determined using NMR (Li *et al.*, 2016). Interestingly, while the superposition of most of the CNBHD (K759-D865) of the calculated ensemble demonstrated a backbone atoms root mean square deviation (RMSD) of 0.49 Å, suggesting a tight and rigid fold in solution, the initial part of the C-linker (R734-P742) displayed high conformational variability (Fig. 2A). Thus, in order to discern whether some of the hKCNH2 CNBHD solution conformations, sampled by the NMR, are shared with its crystal structure, resolved from a tightly packed environment, we next compared the two structures (Fig. 2B). This comparison revealed several conformational differences. Specifically, while the crystal structure clearly indicates that the remnant C-linker maintains it secondary structure and consists of helices α’E and α’F, the solution structure reveals an almost complete unfolding of this region, with a structural pivot observed at G749. Of note, as the C-linker of the NMR sample is preceded by an N-terminal hexa-histidine purification tag and a TEV protease recognition site, due to the reported decreased solubility following proteolysis, the observed flexibility of this region in solution may stem from this non-native sequence addition. Moreover, helices αB and αC show a rigid-body displacement of 8.1 Å (S851-S851; cβ-cβ), resulting from a spatial vacancy due to the complete loss of secondary structure of the α’F helix, associated with the destabilization of the C-linker. Importantly, while the αC-β9 linker is affected by these rearrangements, the ILM itself maintains its spatial localization (Fig. 2B).

**Figure 2.**
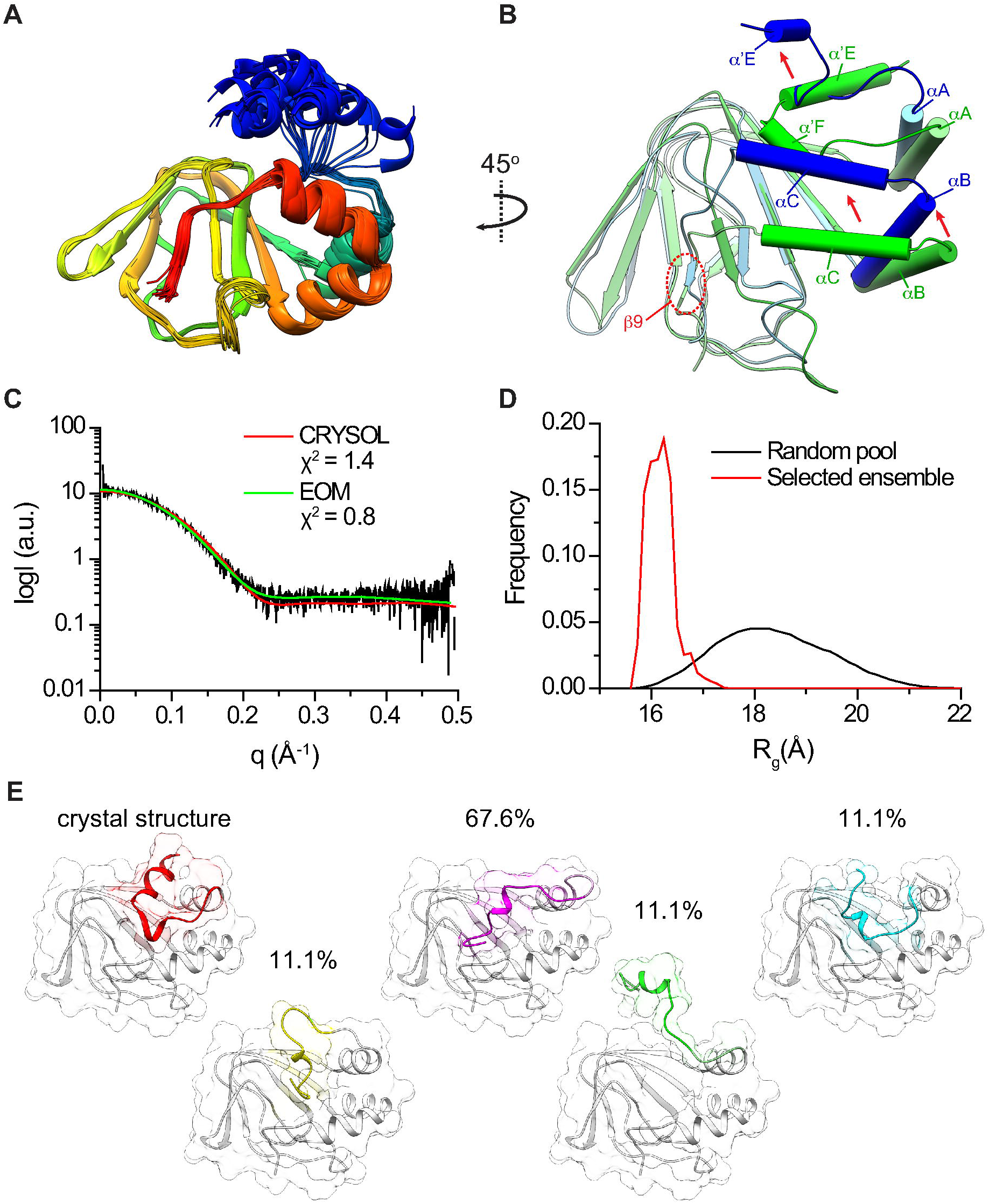
Solution structure of hKCNH2 CNBHD. **(A)** Rainbow colored depiction of the solution NMR structure ensemble (Protein Data Bank code 2N7G) overlay. **(B)** Superposition of the crystal and NMR (ensemble 1) structures of hKCNH2 CNBHD. The β-roll was used as the alignment anchor. **(C)** The experimental SAXS curve of hKCNH2 CNBHD (black curve) was fit using CRYSOL or EOM, as indicated. **(D)** Random R_g_ pool (black curve) and EOM-selected ensemble distribution of the CNBHD (red curve). **(E)** Representative conformations of hKCNH2 CNBHD selected in the EOM analyses. The relative frequency of each conformation in the selected ensemble is indicated. The remnant C-linker domains, acting as the flexible region, are presented in various colors.

To determine whether the structural variability observed stems from the differences in the experimental environments, we proceeded with SAXS analysis of the crystallized hKCNH2 CNBHD construct, subjected to an identical purification scheme. Following data collection, we used CRYSOL (Svergun *et al.*, 1995) to calculate the solution scattering originating from the crystal structure in order to compare it with the experimental SAXS curve. The crystallographic structure demonstrates a reasonable match with the observed solution scattering curve, with χ^2^ = 1.4 (Fig. 2C, Table 2). Next, to explore the possible flexibility of the N-terminus, as observed in the solution NMR structure (Fig. 2A), we employed an ensemble optimization method (EOM) analysis (Bernadó *et al.*, 2007). This approach utilizes an averaged theoretical scattering intensity, derived from an ensemble of conformations, for experimental SAXS data fitting, thus enabling the determination of conformational distributions in solution. First, a random pool of 10,000 conformations was obtained by constructing the residues P742-T747, based on stereochemical restraints. Next, the ensemble of conformations that best fit the data was chosen using a genetic algorithm. Importantly, the selected ensemble fits the data very well (χ^2^ = 0.8; Fig. 2C, Table 2), displaying a narrow distribution of tightly packed conformations (Fig. 2D and E). Thus, the hKCNH2 CNBHD demonstrates an overall high rigidity in solution, occupying a narrow conformational distribution.

### The ILM of hKCNH2

First identified in the structure of the CNBHD from zebrafish KCNH3 (zELK) (Brelidze *et al.*, 2012), all members of the KCNH family share a unique ILM within their canonical CNBHDs (Fig. 1A). In hKCNH2, this short β-strand (β9) consists of the ^860^FNL^862^ tripeptide, forming direct interactions with the cavity of the β-roll (Fig. 1D). While the ILM is rather conserved, several mild sequence differences exist within the KCNH family. Specifically, the first position of the ILM in drKCNH3 (Brelidze *et al.*, 2012), agKCNH2 (Brelidze *et al.*, 2013) and mKCNH1 (Marques-Carvalho *et al.*, 2012) is a tyrosine residue, positioned in an analogous location as the purine ring of cAMP, when bound to HCN2 channels (Zagotta *et al.*, 2003). Furthermore, this tyrosine participates in hydrogen bond formation with either N697 (drKCNH3) or K658 (agKCNH2) (Fig. 1A), suggesting a possible role for its hydroxyl moiety in the spatial organization of the ILM within the β-roll. Conversely, in hKCNH2 this tyrosine is replaced with F860, while the overall fold of the CNBHD is overwhelmingly conserved (Fig. 1D). Thus, a different interaction mechanism must underly the spatial organization of the ILM in hKCNH2. Our structure demonstrates that the position of F860 is governed by a hydrophobic interaction with I789, (5.3 Å; cβ-cβ) and a cation-π interaction with R791, both localized to the β4 strand, and a hydrophobic interaction with L862 of the ILM itself (Fig. 3A). Indeed, the cavity of the CNBHD is devoid of water molecules, supporting the hydrophobic nature of this interaction network. Furthermore, we detected an additional supportive polar interaction network (Fig. 3A). likely playing a crucial role for supporting the ILM conformation and normal channel function (see below). Thus, while structurally conserved, the molecular determinants underlying the spatial positioning of the ILM differ among KCNH family members.

**Figure 3.**
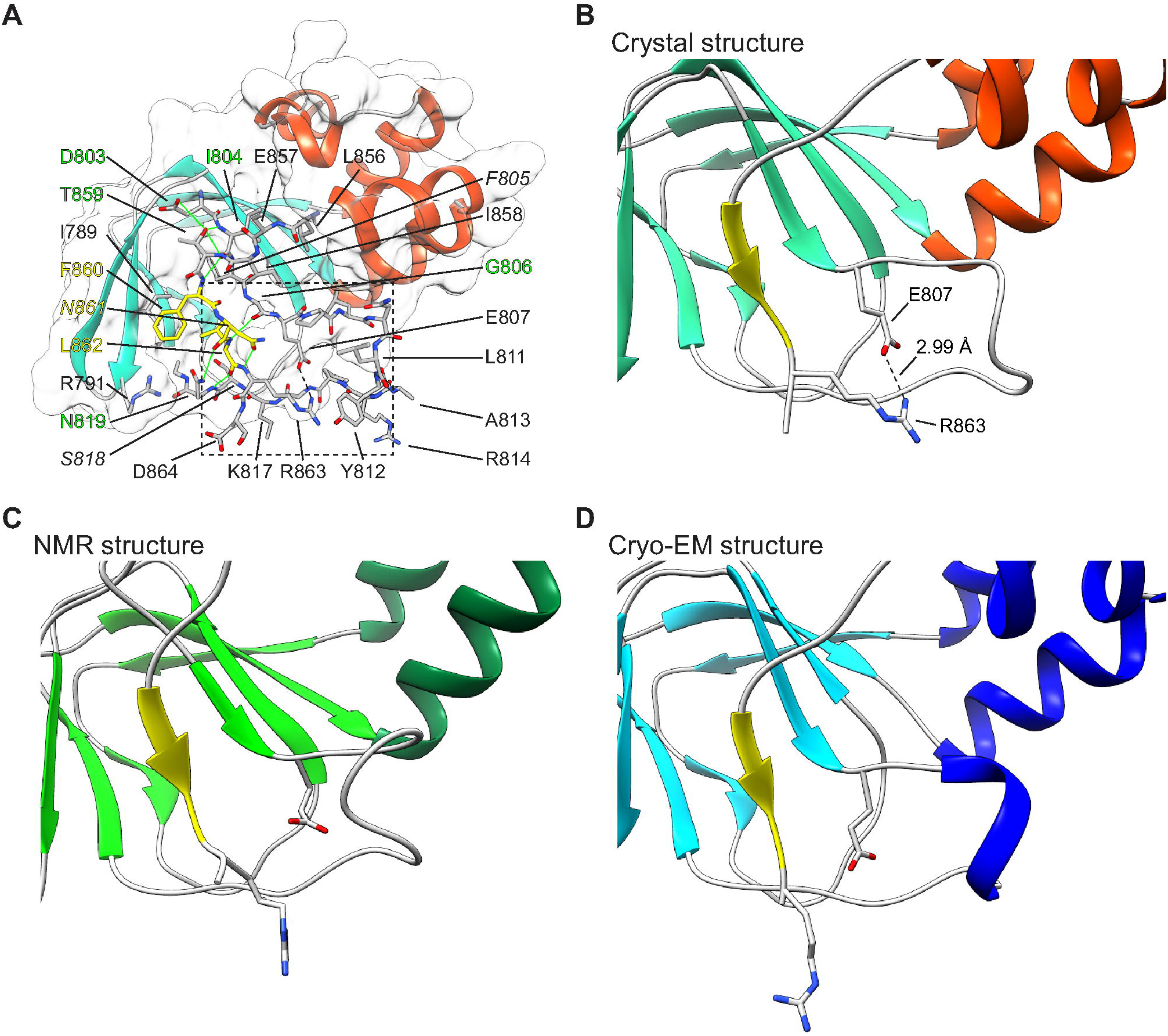
The ILM interaction network. **(A)** Electrostatic surface representation of the CNBHD scaffolding region. Residues which interact with or stabilize the ILM are shown as sticks. ILM, scaffold and hydrogen bond network residues are labeled in yellow, black and green, respectively. LQTS-related residues are highlighted in *italics*. Atoms interacting via hydrogen-bonds and salt-bridge are connected with green and dashed black lines, respectively. **(B-D)** Blowout of the hKCNH2 E807-R863 salt-bridge **(B)**, and the identical region of the NMR (Protein Data Bank code 2N7G) **(C)** and cryo-EM (Protein Data Bank code 5VA1) **(D)** structures. The region shown corresponds to the box indicated in (A).

### An ILM stabilizing salt-bridge is crucial for hKCNH2 channel function

Our structure of the of the CNBHD of hKCNH2 reveals that the ILM is anchored to the β-roll through an intricate network of polar and charged interactions. Indeed, close examination of this region shows that the β9 strand is juxtaposed in parallel of a β-sheet consisting roughly one half of the canonical β-roll (β6-β3-β8; Fig. 1C). This spatial organization is supported by several protein backbone (I804-F860, G806-L862 and N819-L862) and side chain-backbone (T859-I804, R863-N861, N819-N861 and N819-R863) hydrogen-bonds (Fig. 3A). Importantly, additional to these bonds, we detected a single, strategically localized salt-bridge formed between E807 and R863 (Fig. 3B). Indeed, the spatial organization of this salt-bridge suggests it participates in stabilization of the ILM fold. However, this salt-bridge was not observed neither in the NMR structure of the isolated CNBHD (Fig. 3C) nor in the cryo-EM structure of the intact channel (Fig. 3D).

Previously, charge reversal of E807 (E807K) was shown to result in a complete loss of channel function (Cui *et al.*, 2001), indicating the functional importance of the newly identified salt-bridge. To further explore this role, we introduced charge reversal mutations to both positions and tested the properties in Xenopus oocytes using two electrode voltage clamp. Similar to the previously reported results, E807K abolished channel function and no currents could be detected (Fig. 4A). Strikingly, the introduction of an additional charge reversal at the counterpart position (E807K/R863E) resulted in current rescue (Fig. 4A). Analysis of the peak amplitudes of the tail currents, measured in response to a series of step pulses between −80 and 40 mV in 10 mV increments, followed by a subsequent step pulse to −40 mV, demonstrated comparable current amplitudes (Fig. 4A). The G-V relationships further demonstrated WT-like voltage dependency (WT V_50_ = −29.31 ± 0.98 mV, n = 23, E807K-R863E V_50_ = −30.04 ± 0.41, n = 25; Fig. 4C).

**Figure 4.**
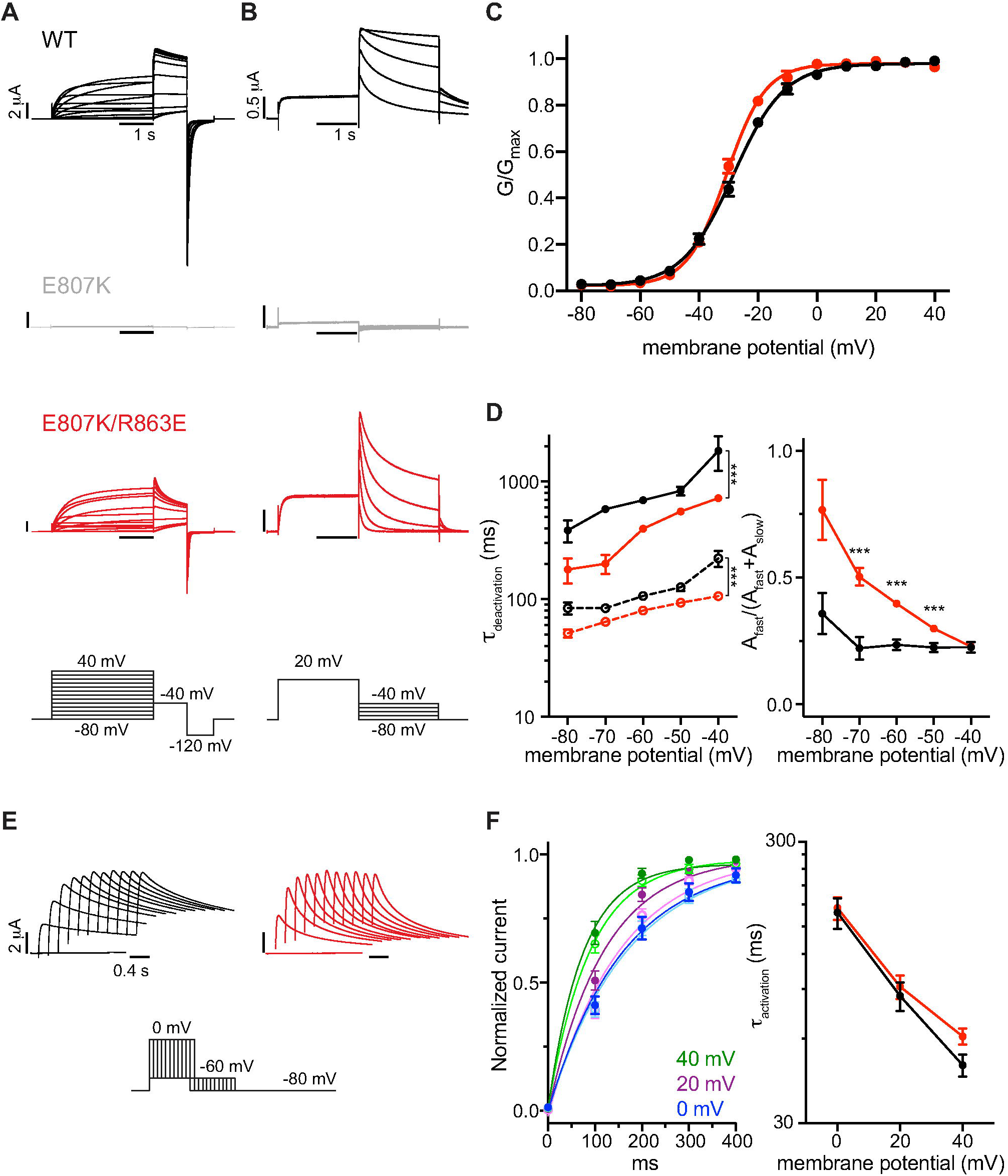
Functional analysis of the E807-R863 salt-bridge. **(A)** Representative current recordings from WT (black), E807K (grey) and E807K/R863E (red). The step pulse protocol is depicted at the bottom. **(B)** Representative tail current recordings for measurement of channels deactivation kinetics. The step pulse protocol is depicted at the bottom. **(C)** G–V relationships. The peak amplitudes of the tail currents at −40 mV were normalized to the maximum tail current in each cell, and the average values are plotted as a function of the membrane potential. The V_50_ values were obtained by fitting the G–V relationships to a Boltzmann function (error bars indicate SEM; n=23-25 cells) **(D)** τ_slow_ and τ_fast_ values of deactivation represented as solid and dashed lines, respectively (left). Deactivation time constants for WT (black) and E807K/R863E (red). Each deactivating tail current was fit to a double exponential function to calculate the slow and fast time constants, and the average values were plotted as a function of membrane potential. Fractional fast component contribution, A_fast_/(A_fast_+A_slow_), is presented as a function of membrane potential (right) (error bars indicate SEM; n=11-17 cells; *** *p* < 0.005, Student’s t-test). **(E)** Representative activation current recordings from WT (black) and E807K/R863E (red). Varying durations of depolarizing test pulses were followed by a step pulse to −60 mV, as shown at the bottom. **(F)** The activation kinetics were evaluated by the monoexponential fit of the increase of the peak tail current amplitude in response to incremental increase in test pulse duration (left). WT and E807K/R863E activation curves are shown as closed and open symbols, respectively, colored according to the test pulse voltage used. Evaluated τ_activation_ values at 0, 20 and 40 mV of WT (black) and E807K/R863E (red) are presented (right).

Next, we investigated the deactivation kinetics of the double reversal mutant. Deactivating current traces of WT and mutant were obtained by a step activation to 20 mV, followed by deactivation pulses ranging from −40 to −80 mV (Fig. 4B). To determine the slow and fast time constants, tail currents were fit with a double exponential function (τ_slow_ and τ_fast_, respectively). Intriguingly, although showing an overall similar voltage dependency (Fig. 4C), the double mutant displays significantly accelerated deactivation kinetics, compared with the characteristic slow deactivation of the WT channel (Fig. 4D). Specifically, while the WT channel deactivation followed a relatively slow time course (τ_slow_ = 694 ± 33 ms and τ_fast_ = 106 ± 6 ms at −60 mV, n = 11), the double mutant demonstrated faster kinetics (τ_slow_ = 200 ± 36 ms and τ_fast_ = 80 ± 2 ms at −60 mV, n = 17; *p* < 0.001 for both time constants). Moreover, analysis of the fractional contribution of the fast and slow components showed that while the WT channels demonstrate a voltage insensitive balance, leaning towards the slow component, the fast component dominates the double mutant kinetics at more hyperpolarized potentials (Fig. 4D).

The salt-bridge charge reversal mutations resulted in faster channel deactivation kinetics (Fig. 4D), while keeping voltage dependency unaffected (Fig. 4C). A possible gating mechanism accommodating for these two effects may involve activation kinetics acceleration. Indeed, perturbation of the only inter-domain salt-bridge observed in the mouse KCNH1 eag domain-CNBHD complex structure (R57-D642; Fig. 5A) resulted in acceleration of channel activation (Haitin *et al.*, 2013). Thus, to discern whether a similar mechanism is present here, we analyzed the activation kinetics of WT and E807K/R863E by measuring the tail current amplitude dependence on the increase of the preceding test potentials duration (Fig. 4E). Next, a mono-exponential function was employed to fit the activation time course and obtain the τ_activation_ time constant. Intriguingly, the activation kinetics of E807K/R863E were not significantly changed throughout the voltage clamp regimes tested (Fig. 4F; n = 6-9; *p* > 0.05). Thus, activation kinetics are unaffected by the charge reversal mutations. Together, the crystal structure of hKCNH2 CNBHD reveals a novel functionally important E807-R863 salt-bridge, which plays a role in channel deactivation.

**Figure 5.**
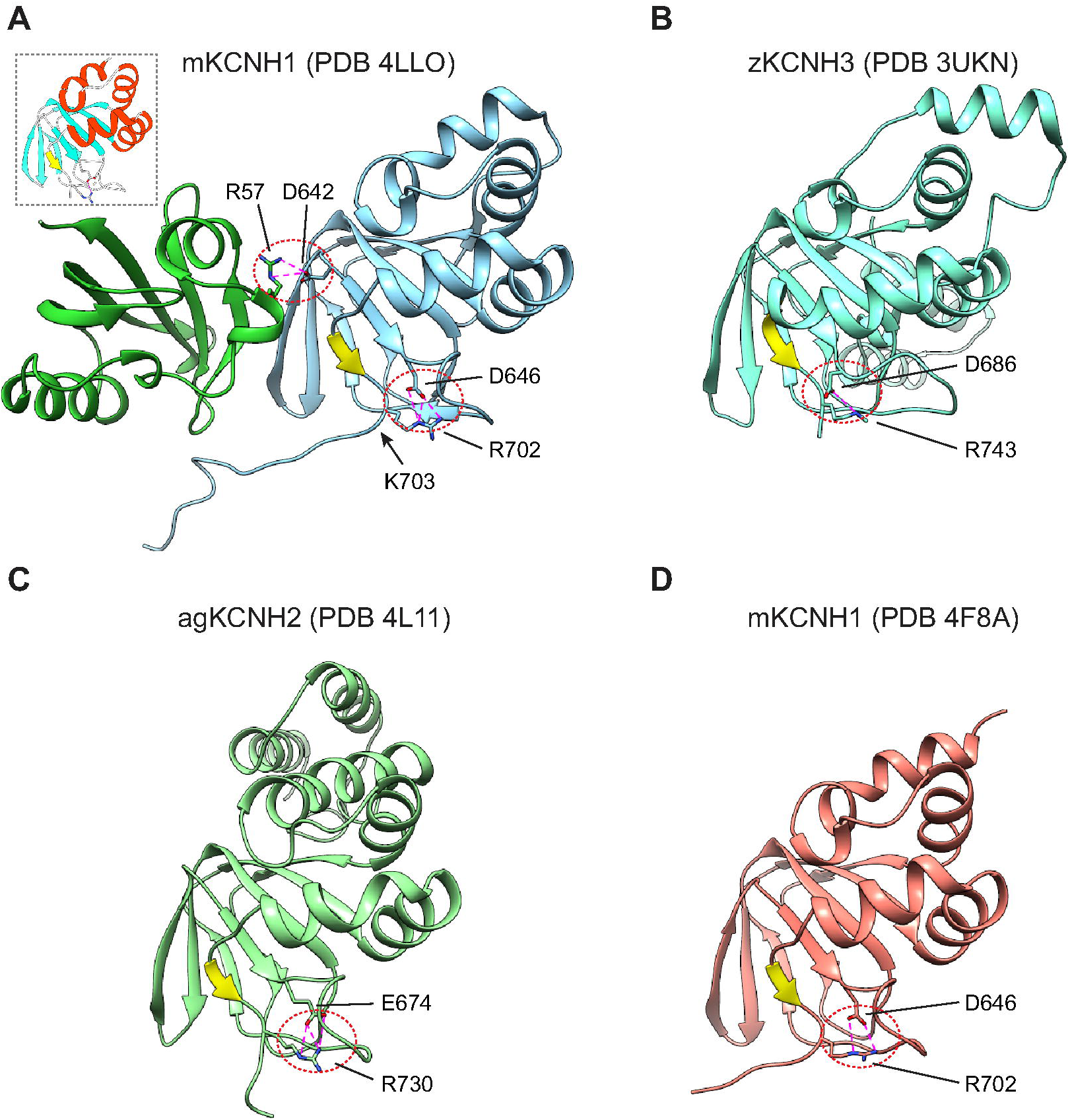
Structural comparisons with select KCNH CNBHD crystal structures. **(A)** Structure of the eag domain-CNBHD complex from mouse KCNH1. The N-terminal eag domain is colored green, the C-terminal CNBHD is colored blue. Inter-domain (R57-D642) and CNBHD (D646-R702) salt-bridges are indicated in magenta. Arrow indicates the post-CNBHD kink. G749 The CNBHD of hKCNH2 is presented for orientation (inset). **(B-D)** Salt-bridges of the CNBHDs of zKCNH3 (D686-R743) (B), agKCNH2 (E674-R730) (C) and mKCNH1 (D) are indicated as in (A). ILMs are indicated in yellow throughout.

## Discussion

In this study, we have used various biophysical approaches to resolve the molecular determinants involved in the structural organization of hKCNH2 CNBHD. Following the initial characterization of the purified CNBHD using SEC-MALS (Fig. 1B), we crystallized and solved the structure of this domain at high resolution. This structure, solved at an unprecedented resolution for a CNBHD of 1.5 Å (Brelidze *et al.*, 2012, 2013; Marques-Carvalho *et al.*, 2012; Haitin *et al.*, 2013), provided exquisite molecular details into the spatial organization of the ILM (Fig. 1D). Importantly, the crystallographic structure is in high agreement with the solution scattering profile (Fig. 2C-E) and the recent NMR analysis of hKCNH2 (Li *et al.*, 2016). Importantly, the atomic structure resolution identified a novel salt-bridge, which was not previously resolved (Fig. 3). Strikingly, electrophysiological analysis of charge reversal mutations revealed its crucial role for hKCNH2 function (Fig. 4).

KCNH channels are widely expressed in the human body and play central roles in numerous physiological and pathophysiological processes. hKCNH2 channels, arguably the prominent pathophysiologically-relevant family members, have a critical role in human cardiac repolarization. Unlike the cyclic-nucleotide gated (CNG) and HCN channels, the CNBHD of KCNH encompasses the ILM, spatially replacing the cyclic nucleotides, blocking their binding site (Brelidze *et al.*, 2012; Marques-Carvalho *et al.*, 2012; Haitin *et al.*, 2013). Ever since its identification (Brelidze *et al.*, 2012; Marques-Carvalho *et al.*, 2012), the ILM was shown to play a crucial role in KCNH channel family function (Brelidze *et al.*, 2013; Zhao *et al.*, 2017; Codding & Trudeau, 2019). Nevertheless, the mechanism by which it participates in channel gating modulation remains ambiguous. Indeed, possible modulatory mechanisms may involve the interaction of the CNBHD with the canonical N-terminal eag domain (Gianulis *et al.*, 2013; Haitin *et al.*, 2013), together with alterations of the conformational space occupied by the CNBHD and its associated C-linker (Zhao *et al.*, 2017).

Over that past years, major strides forward have been made towards a comprehensive elucidation of the structure-function relations governing KCNH channels gating, culminating in the determination of the structure of an intact hKCNH2 channel by cryo-EM (Wang & MacKinnon, 2017). This groundbreaking milestone provided the long-sought molecular characterization of hKCNH2 transmembrane domain, depicting the voltage sensors in an active conformation, explaining their notorious sensitivity to drug-induced blockade and shedding light on the fast inactivation mechanism. Moreover, the spatial organization of the intracellular eag domain and CNBHD was reported. However, due to the inherent limitations of cryo-EM, this functionally important intracellular region was resolved at lower resolution. Therefore, the exact interaction network underlying CNBHD occupancy by the ILM in hKCNH2 remained unanswered.

Previous studies focused on the ILM itself and its effect on the interaction between the CNBHD and the eag domain (Zhao *et al.*, 2017; Codding & Trudeau, 2019). Here, based on our structural analysis, we focused on a putative salt-bridge which deemed important for stabilization of the ILM. Structural comparison of our structure with previously determined crystal structures of KCNH CNBHD homologs revealed that the spatial organization leading formation of the E807-R863 salt-bridge is highly conserved (Fig. 5). Indeed, CNBHD structures from zKCNH3 (Fig. 5B), agKCNH2 (Fig. 5C) and mKCNH1 (Fig. 5D) show that positions homologous to E807 and R863 of hKCNH2 are nearly identically positioned. As shown here, consistent with the high level of structural conservation, the salt-bridge plays a role in channel function (Fig. 4). Previously, based on electrophysiological and spectroscopic studies of intact channels, it was suggested that the interaction between the CNBHD and the eag domain plays a role in the functional regulation of KCNH channels (Dai & Zagotta, 2017; Codding & Trudeau, 2019). However, the only available high-resolution structure of the intracellular complex of KCNH channels remains the co-crystal structure of the mouse KCNH1 eag domain-CNBHD (Haitin *et al.*, 2013). The structural alignment revealed a kink immediately following the conserved salt-bridge (K703), with the post-CNBHD region, encompassing a calmodulin binding site, cradling the eag domain (Fig. 5A). Thus, it appears that the salt-bridge serves as a linchpin, limiting the conformational landscape of the ILM, while allowing interaction of the post-CNBHD with the eag domain in mouse KCNH1 channels. This can support recent results demonstrating that the ILM undergoes a defined conformational change during voltage-dependent channel potentiation, rather than dissociation from its binding pocket upon the relative rearrangement between the eag domain and the CNHBD (Dai *et al.*, 2018).

In conclusion, this study provides molecular insights into the structural organization of the hKCNH2 CNBHD in an unprecedented resolution. This structure revealed a previously overlooked salt-bridge, playing a role in channel function. Structural comparison with the intracellular complex of mouse KCNH1, together with electrophysiological studies, suggests that this salt-bridge is strategically positioned to support both the organization of the ILM and the intracellular complex interface. As most KCNH channels differ in their post-CNBHD region, future studies are needed in order to address the functional role of this salt-bridge during channel gating and regulation in different family members.

## Acknowledgements

We thank the staff of beamlines ID23-1 and BM29 at the ESRF for assistance with diffraction experimentation. We also thank Dr. Moran Rubinstein and members of the Haitin lab for advice and support. This work was performed in partial fulfillment of the requirements for a Ph.D. degree of A.B., Sackler Faculty of Medicine, Tel Aviv University, Israel. This work was supported by the Israel Science Foundation (grant number 1721/16), the Israel Cancer Research Foundation (grant number 01214) and from the German-Israeli Foundation (grant number I-2425-418.13/2016). Supported also came from the I-CORE Program of the Planning and Budgeting Committee and The Israel Science Foundation (grant number 1775/12).

## Author contributions

Conceived and designed the experiments: Ariel Ben-Bassat and Yoni Haitin; Performed the experiments: Ariel Ben-Bassat; Analyzed the data: Ariel Ben-Bassat and Moshe Giladi; Wrote the paper Ariel Ben-Bassat, Moshe Giladi and Yoni Haitin.

## Accession Number

Atomic coordinates and structure factors have been deposited in the Protein Data Bank with accession number 6SYG.

